# Intergenerational transmission of education and ADHD: Effects of parental genotypes

**DOI:** 10.1101/664128

**Authors:** Eveline L. de Zeeuw, Jouke-Jan Hottenga, Klaasjan G. Ouwens, Conor V. Dolan, Erik A. Ehli, Gareth E. Davies, Dorret I. Boomsma, Elsje van Bergen

**Affiliations:** Department of Biological Psychology, Vrije Universiteit, Amsterdam, the Netherlands; Amsterdam Public Health Research Institute, VUmc, Amsterdam, the Netherlands; Avera Institute for Human Genetics, Avera McKennan Hospital & University Health Center, Sioux Falls, SD, US

**Keywords:** Intergenerational Transmission, Genetic Nurturing, Polygenic Scores, Educational Attainment, Academic Achievement, ADHD

## Abstract

It is challenging to study whether children resemble their parents due to nature, nurture, or a mixture of both. Here we used a novel design that employs the fact that parents transmit 50% of their alleles to their offspring. The combined effect of these transmitted and non-transmitted alleles on a trait are summarized in a polygenic score (PGS). The non-transmitted PGS can only affect offspring through the environment, via genetically influenced behaviours in the parents, called genetic nurturing. For genotyped mother-father-offspring trios (1,120-2,518 per analysis) we calculated transmitted and non-transmitted PGSs for adult educational attainment (EA) and childhood ADHD and tested if these predicted outcomes in offspring. In adults, both transmitted (R^2^ = 7.6%) and non-transmitted (R^2^ = 1.7%) EA PGSs predicted offspring EA, evidencing genetic nurturing. In children around age 12, academic achievement was predicted only by transmitted EA PGSs (R^2^ = 5.7%), but we did not find genetic nurturing (R^2^ ∼ 0.1%). The ADHD PGSs did not significantly predict academic achievement (R^2^ ∼ 0.6%). ADHD symptoms in children were predicted by transmitted EA PGSs and ADHD PGSs (R^2^ = 1-2%). Based on these results, we conclude that previously reported associations between parent characteristics and offspring outcomes seem to be mainly a marker of genetic effects shared by parents and children.

## Introduction

Research on influences of the family environment on children’s behaviour is hampered by the fact that parents provide their offspring with both the family environment and genes, leading to intertwined effects (Plomin et al. 1977). The resulting gene-environment correlation arises from environmental experiences that are correlated with genetic propensities. Gene-environment correlations can be 1) passive, in which the rearing environments that parents create are related to the parents’ and hence to the children’s genotypes, 2) evocative, in which children evoke responses from others that are related to their genotype, and 3) active, in which children actively seek out environments that are related to their genotypes. Passive gene-environment correlations may explain well-established associations between home characteristics and child outcomes. Examples thereof are, the association between the number of books in the home and children’s reading skills (van Bergen et al. 2018; Sikora et al. 2019) and the association between household chaos and children’s problem behaviour (Coldwell et al. 2006). What is needed is a design that controls for children’s genetic propensities to demonstrate genuine family environmental influences. Here we study children’s academic achievement and ADHD symptoms and seek evidence for influences passed on from parent to child through the rearing environment, utilizing the family members’ measured DNA.

Last year two research groups independently developed a novel design that employs the genotypes of a child and his/her parents to demonstrate the influence of the child’s heritable rearing environment (Bates et al. 2018; Kong et al. 2018). At each autosomal locus in the human genome, parents transmit one of the two homologous alleles to their offspring. In genotyped parents and offspring trios it can be determined which alleles were transmitted and which were not. In such offspring, one can calculate two polygenic scores (PGS): one based on the transmitted alleles and one based on non-transmitted alleles. The transmitted PGS for educational attainment (EA) aggregates thousands of small genetic effects that together explain ∼12% of variance in people’s EA (Lee et al. 2018). Bates et al. and Kong et al. demonstrated, as expected, that the EA PGS of the transmitted alleles explained variance in offspring EA in (young) adulthood. But strikingly, the EA PGS based on the non-transmitted alleles of their parents also explained variance in offspring EA. As only the parents, not the offspring, carry these alleles, this effect must be due to an indirect effect of the parental genotype on the EA of the offspring. A plausible explanation is that the parental contribution to the offspring environment (relevant to EA) is in part a function of their genotype. Hence, Kong et al. termed this phenomenon ‘genetic nurturing’. Since the transmitted alleles are present in both the parents and the offspring, the transmitted alleles might have a direct effect on offspring’s behaviour, but they can also have an effect through the environment that parents create based on their EA. Therefore, transmitted alleles induce both genetic nurturing and direct genetic effects while non-transmitted alleles only induce genetic nurturing.

In the current study, we applied this transmitted and non-transmitted PGS design to seek proof for influences of the family environment on two important and heritable childhood traits, academic achievement and ADHD. In a classical twin design, genetic nurturing would be captured in the common environmental variance, because un-modelled gene-environment correlation contributes to this type of variance in a classical twin design (Purcell 2002), and enetic nurturing is a ‘true’ effect of the home environment that is shared between twins (Posthuma et al. 2003). Genetic nurturing, if present, should also be reflected by the presence of environmental transmission in extended family studies, such as adoption, parent-offspring and children-of-twins designs, as these designs can separate parent-offspring resemblance into a part due to the family environment and a part due to genetic transmission (D’Onofrio et al. 2003). In studies utilizing a PGS from one of the parents, if a maternal or paternal PGS significantly predicts trait differences in offspring, even after correcting for the effect of a child’s own PGS, this would too point to genetic nurturing (Belsky et al. 2018).

For academic achievement and ADHD, we had opposing hypotheses regarding genetic nurturing based on previous findings from behaviour genetic studies. Academic achievement in primary school is substantially heritable (∼75%) with modest common environmental influences (∼10%) (de Zeeuw et al. 2016). Associations between parental EA and academic achievement in adopted children have been found in some, but not all, adoption studies (Lundborg et al. 2018). In addition, a parent-offspring study of EA in adulthood found both genetic and environmental transmission (McGue et al. 2017). Children-of-twins studies also indicate that intergenerational transmission of education is, at least partly, explained by an effect of parental EA that goes through the home environment (e.g. Chevalier et al. 2013). Finally, a maternal PGS for EA predicted a child’s performance at the end of secondary school beyond the effect of its own EA PGS, and this effect was mediated by parenting behaviours (Wertz 2019). Based on these findings we expected a small role of genetic nurturing in academic achievement in children. Heritability of ADHD is also high (∼75%) in both case-control and general population samples (Faraone and Larsson 2019), without evidence for the influence of the family environment. There are even indications that dominant genetic effects play a role (Rietveld et al. 2003; Nikolas and Burt 2010), which would hamper the detection of common environmental effects in a classical twin design. The presence of common environmental variance is not actually ruled out when a model including dominant genetic effects fits the twin data well. Extended family studies focusing on the transmission of ADHD within families are scarce (McAdams et al. 2014). A study of twin families with adult twins and their siblings in the offspring generation established that familial transmission was explained by genetic inheritance, without support for cultural transmission from parents to offspring (Boomsma et al. 2010). Also, according to a children-of-twins study, the link between antisocial behaviour in parents and ADHD in offspring is solely due to a transmission of a genetic liability (Silberg et al. 2012). Based on these previous studies we did not, in contrast to for academic achievement, expect to find an effect of genetic nurturing on offspring ADHD symptoms.

In this study, we first aimed to replicate the finding of an effect of genetic nurturing of EA on adult offspring EA from Kong et al. (2018) and Bates et al. (2018) in a Dutch sample. Subsequently, we investigated the presence of genetic nurturing on childhood academic achievement and ADHD symptoms. We considered two continuous measures of ADHD symptoms based on maternal and teacher ratings, as rater agreement between mother and teacher is relatively low (*r* = .44) (Achenbach & Rescorla, 2001), possibly suggesting that children behave differently in a home versus a school situation. We examined the effect of transmitted and non-transmitted parental alleles, quantified in terms of PGSs, on academic achievement and ADHD symptoms at the end of primary school. As ADHD symptoms are related to academic achievement (Polderman et al. 2010), we tested effects both within developmental domains, and across domains, with and without taking the effects within domain into account. In sum, we investigated whether the transmitted PGS and non-transmitted PGS of 1) EA predicts adults EA, 2) EA predicts children’s academic achievement, 3) ADHD predicts children’s ADHD symptoms, 4) EA predicts children’s ADHD symptoms and 5) ADHD predicts children’s academic achievement.

## Methods

### Participants

The Netherlands Twin Register (NTR) was established around 1987 by the Department of Biological Psychology at the Vrije Universiteit Amsterdam and recruits approximately 40 per cent of new-born twins or higher-order multiples in the Netherlands for longitudinal research. Parents of twins receive a survey about the development of their children every two to three years until the twins are 12 years old. Starting at age 7 years, parents are asked consent to also approach the primary school teacher(s) of their twin and other children. The survey sent to mothers, fathers and teachers includes the age and context appropriate version of the Achenbach System of Empirically Based Assessment (ASEBA) (Achenbach, Ivanova, & Rescorla, 2017). Adult twins were registered with the NTR through several approaches, including, for example, recruitment through city council offices in the Netherlands, advertising in NTR newsletters and the internet. Parents, siblings, spouses and offspring of adult twins are also invited to take part. Since 1991, participants receive a survey every two to three years with questions on, amongst others, health, personality, and lifestyle. The NTR has also been collecting genotyping data in both children and adults in several large projects. More details concerning the NTR’s data collection, the methods of recruitment, participants’ background and response rates are described elsewhere (van Beijsterveldt et al. 2013; Willemsen et al. 2013). The data collection was approved by the medical ethical review committee of the VU Medical Center Amsterdam (NTR25052007) and informed consent was obtained from all participants included in the study.

For 5,900 offspring (from 2,649 families) their own, as well as the genotype data of both of their parents were available. Data were excluded if an individual had a non-European ancestry (n = 472). In this group of families, information on EA was available for 1,931 adult offspring (662 males and 1,260 females) from birth cohorts 1946-1991 (Age at assessment of EA: Mean = 33.9, SD = 7.51, Range = 25-64 year). Childhood academic achievement scores were available for 1,120 offspring around age 12 (509 boys and 611 girls) from birth cohorts 1983-2002. Data on ADHD symptoms at home were available for 2,518 children (1,202 boys and 1,316 girls) from birth cohorts 1986-2008. Data on ADHD symptoms at school were available for 1,969 (968 boys and 1,001 girls) children from birth cohorts 1986-2011

### Measures

#### Educational attainment

EA in adults was measured by means of a self-report on highest obtained degree. The responses were recoded into four categories: primary education (level 0), secondary education (level 1), higher secondary education (level 2) and tertiary education (level 3). EA was only analyzed in individuals over 25 years of age.

#### Academic achievement

Academic achievement in children was assessed around age 12 by a nationwide standardized educational achievement test (Cito 2002). The results on this test are, in combination with teacher advice, used to determine the most suitable level of secondary education. Around 75% (before 2015) or 50% (after 2015) of Dutch children took this test in their final year of primary school. The test consists of multiple choice items in four domains, namely Arithmetic, Language, Study Skills and Science and Social Studies. The first three test scales are combined into a Total Score, which is converted into a score ranging from 500 to 550, which reflects the child’s standing relative to the total group of children who took the test in a given school year (e.g. in 2015 this group comprised ∼165,000 children) (van Boxtel et al. 2010).

#### ADHD symptoms

ADHD symptoms were assessed with the ASEBA system empirically based syndrome Attention Problems scale (Achenbach et al. 2017). The Child Behavior Check List (CBCL) for school aged children (6-18 years) was used to assess behaviour at home, and the Teacher Report Form (TRF) for behaviour at school The ASEBA Attention Problems scale includes items on inattention (e.g. ‘Fails to finish things he/she starts’; CBCL: 2 items and TRF: 13). and hyperactivity/impulsivity (e.g. ‘Can’t sit still’; CBCL: 2 items and TRF: 10) The items are scored on a 3 point scale from 0 (‘not true or never’) to 2 (‘completely true or very often’). Missing items were imputed by the average item score of the scale for a child. Missing items were imputed by the average item score of the scale for a child, if missingness on the scale items was less than 20%. ADHD symptoms were assessed at age 10 and 12, and, if available, age 12 data were analysed, or else age 10 data. The data showed an L-shaped distribution and were square root transformed prior to analyses.

#### Genotyping

Genotyping and data QC in a twin register involves dealing with (nearly) identical DNA of monozygotic twins and triplets. In our cohort, genotyping is used for determining zygosity as described earlier (Odintsova et al. 2018) as well as multiple scientific analyses. While the former needs both twins of a pair to be genotyped, for the latter this is not necessarily true, and for several steps in genotype data QC and statistical analyses having both twins in the data can even introduce problems if not taken into account (Minică et al. 2017). Hence, if the DNA data of monozygotic twins and triplets is indeed DNA confirmed identical, a single sample can be selected to run several steps of the data QC, and subsequently the DNA of this twin is duplicated back to the other twin at the most convenient moment as described below.

The genotype data used for this study included 17,620 unique DNA samples, done on several different platforms: Affymetrix-Perlegen (n = 1,117), Illumina 660 (n = 1,323), Illumina Omni Express 1 M (n = 234), Affymetrix 6.0 (n = 7,086), Affymetrix Axiom (n = 2,665) and Illumina GSA (n = 5,195). Genotype calls were made with the platform specific software (Birdseed, APT-Genotyper, Beadstudio) following manufacturers’ protocols. For the Affymetrix-Perlegen and Illumina 660 platforms, the single nucleotide polymorphisms (SNPs) were lifted over to build 37 (HG19) of the Human reference genome.

Per platform, a sample was removed if the call rate for this person was < 90%, the Plink 1.07 F heterozygosity value was < −0.10 or > 0.10, the gender of the person did not match the DNA of the person, the IBD status did not match the expected familial relations, or the sample had > 20 Mendelian errors. In case a subject was genotyped on multiple platforms, only the platform with the highest number of SNPs was selected if genotypes were concordant (> 0.97). Allele and strand alignment of SNPs was done against the Dutch Genome of the Netherlands (GONL) reference panel for each platform (Boomsma et al. 2014). SNPs were removed in each platform when Minor Allele Frequency (MAF) < 0.01, Hardy-Weinberg Equilibrium (HWE) test p-value < 10^-5^ or the call rate of the SNP was < 95%. Subsequently, only SNPs were selected if the allele frequency of the SNP deviated < 0.10 as compared to the GONL data. All palindromic SNPs with a MAF > 0.40 were also removed. The individual platform data were then merged into a single dataset. In this dataset, the sample IBD, on a common backbone of ∼70K SNPs, was re-compared with their expected familial relations and samples were removed if these did not match. Subsequently, the data were phased and imputed with Mach-admix, using GONL as a reference panel, for those SNPs that survived quality control and were present on at least one platform, forcing missing genotype imputation for all SNPs. Best guess genotypes were generated from these data, and the following SNPs were selected: SNPs with a R^2^ > 0.90, with HWE p > 10^-5^, with a Mendelian error rate < 2%, and if the association of one platform=case vs. the other platforms=controls p-value > 10^-5^ (applied for each platform) resulting in a genetic backbone of 1.2M SNPs. After this step, 3017 DNA confirmed monozygotic twin samples were returned into the dataset by duplicating the SNP data of their co-twin. Another 364 DNA samples were added, 335 out of the original 349 samples, plus 29 of their confirmed monozygotic twins, of the NTR that were also sequenced in the GONL reference population. The resulting was a final dataset of 21,001 individuals from 6,671 families with 1.2M SNPs. The cross-chip imputed data were used to calculate genetic principal components using SmartPCA software, and the PCAs were subsequently used to determine if a person was from non-European descent (Galinsky et al. 2016). The full set was then aligned against the 1000 genomes phase 3 version 5 reference panel, and imputed on the Michigan imputation server (Das et al. 2016). From the imputed 1000G VCF files, best guess genotypes were calculated for all markers using Plink 1.96.

#### Non-transmitted genotypes

In total there were 2,649 families having two genotyped parents, with 5,900 offspring, including 1,245 MZ twin pairs (Mean = 2, Range = 1-10), for which allele transmission could be calculated on the 1000G imputed data. Before this calculation, the genotype data were filtered using the following criteria: only ACGT SN Ps on the autosomes, no SNPs with duplicate positions, no SNPs with 3 or more alleles, MAF > 0.01, HWE p > 10^-5^ and genotype call rate > 0.99, leaving 7,411,699 SNPs. For the 5,900 offspring, this is the transmitted alleles dataset. Subsequently, all children were defined as being a case, and then the Plink --tucc option was used to generate a single TDT pseudo-control genotype for each child (given the 2 parents), resulting in the non-transmitted alleles dataset. Both datasets were then used to calculate PGSs.

#### Polygenic scores

For the EA PGS calculation we used the GWA summary statistics from the EA meta-analysis (Lee et al. 2018) and for ADHD we used the statistics from the meta-analysis for ADHD (Demontis et al. 2019), both excluding the NTR and 23andMe cohorts. After excluding these cohorts, the meta-analysis was redone for EA and ADHD symptoms. Since the NTR was present in the quantitative EAGLE summary statistics, which were combined with the Psychiatric Genetics Consortium (PGC) case-control ADHD summary statistics, we also re-applied the correction method to join case-control and quantitative summary statistics (Demontis et al. 2019).

Based on these summary statistics sets, linkage disequilibrium (LD) weighted Beta’s were calculated using the LDpred package with the summary statistics top proportion of 0.3 of SNPs (Vilhjálmsson et al. 2015) with an LD pruning window of 250 KB. The reference population to calculate the LD patterns was a selection of the first 2500 2^nd^ degree unrelated 1000G imputed individuals from the 5,900 NTR individuals that were used for scoring. The detection of unrelated individuals was done with the King software (Manichaikul et al. 2010). The resulting LD corrected Beta’s were used to calculate polygenic scores using the Plink 1.90 software, in the transmitted and non-transmitted alleles datasets.

### Statistical Analyses

The current study included one offspring outcome in adulthood, i.e. EA, and two in childhood, i.e. academic achievement and ADHD symptoms. In adulthood, EA was regressed on the transmitted and non-transmitted EA PGS to replicate previous findings (Bates et al. 2018; Kong et al. 2018). In childhood, academic achievement, ADHD symptoms at home and ADHD symptoms at school were regressed on the transmitted and non-transmitted EA PGSs (model 1), on the transmitted and non-transmitted ADHD PGSs (model 2) and on the transmitted and non-transmitted EA plus ADHD PGSs (model 3). All outcome measures were residualized for the effects of sex, year of birth (only for EA), the interaction between sex and year of birth (only for EA), 10 principal components reflecting Dutch ancestry differences, and the genotyping platform. Within each analysis, the predictors and residualized outcome measures were standardized in the subset of individuals that had both PGS and phenotype data. A random intercept was added to correct for dependency of the observations due to family clustering. Generalized linear models were fitted in the statistical program SPSS Statistics for Windows 25.0 (IBM Corp., 2017) with maximum likelihood estimation. The type of model depended on the measurement level of the outcome: EA (ordinal logistic), academic achievement (linear) and ADHD symptoms (linear). To correct for multiple testing an alpha level of 0.01 was adopted.

### Power analysis

The sample included twin pairs and their siblings, which meant that observations were not independent. To facilitate the power analysis, we used the effective sample size, i.e. N_E_ = (N*M)/(1+ICC*(M-1)) in which N = the number of families, M = the number of individuals in a family and ICC = the (average) phenotypic correlation within a family. We applied this separately for MZ and DZ (and siblings) families, given the expected differences in ICC. The power to detect a particular effect size (i.e. percentage of phenotypic variance explained) of the non-transmitted PGS was based on the non-central F-distribution. Power equals the percentage of significant tests of the regression coefficient given an alpha level of 0.01

## Results

The EA and ADHD PGSs correlated moderately for both the transmitted (*r* = -.271) and non-transmitted (*r* = -.234) PGSs, indicating that genes involved in EA and ADHD partly overlap. The correlation between the transmitted and non-transmitted PGSs was low for EA (*r* = .090) and ADHD (*r* = .033). PGSs and phenotype data were available for adult EA (N = 1,931, level 0 = 0.5%, level 1 = 8.6%, level 2 = 32.0 %, level 3 = 59.0%), and childhood academic achievement (N = 1,120, Mean = 538.8, SD = 8.1, Range = 507-550), ADHD symptoms at home (N = 2,518, Mean = 2.92, SD = 3.2, Range = 0-18) and ADHD symptoms at school (N = 1,969, Mean = 6.09, SD = 7.7, Range = 0-43).

In adulthood, the transmitted (*β* = .28, *p* < .001) and non-transmitted (*β* = .13, *p* < .001) PGSs based on the EA meta-analysis (Lee et al. 2018) significantly predicted EA (see Figure 1A), replicating previous findings for genetic nurturing in EA (Bates et al. 2018; Kong et al. 2018). The magnitude of the estimated effect of the non-transmitted alleles for EA on adult EA was almost half of the effect of the transmitted alleles. In childhood, the transmitted EA PGS significantly predicted academic achievement (*β* = .24, *p* < .001), ADHD symptoms at home (*β* = -.13, *p* < .001) and ADHD symptoms at school (*β* = -.13, *p* < .001) (Table I - model 1). The non-transmitted EA PGS did not have a significant effect (see Figure 1B). This suggests that there was no genetic nurturing on offspring’s academic achievement and ADHD symptoms elicited by parental EA. The transmitted ADHD PGS based on the ADHD meta-analysis (Demontis et al. 2019) did not predict academic achievement (*β* = -.08, *p* = .022), but had a significant influence on ADHD symptoms at home (*β* = .17, *p* < .001) and ADHD symptoms at school (*β* = .13, *p* < .001) (Table I - model 2). The non-transmitted ADHD PGS did not predict any of the outcomes, indicating that the environment that parents provided to their children based on the genes that play a role in their own ADHD symptoms did not affect their children’s development (see Figure 1B). When taking the effects of both the EA and ADHD PGSs into account, the effect of the transmitted EA PGS (*β* = .23, *p* < .001) on academic attainment was similar, but the effect of the ADHD PGS (β = -.02, p = .525) was attenuated. The effects of the transmitted EA PGS (*β* = -.09, *p* < .001) and ADHD PGS (β = .14, p < .001) on ADHD symptoms at home were both diminished. The effects of the transmitted EA PGS (*β* = -.10, *p* < .001) and ADHD PGS (β = .10, p < .001) on ADHD symptoms at home are also somewhat lower (Table I - model 3). See figure 3 for a schematic representation of the results.

**Table I.** The estimated effects (with 95% CI) of the transmitted (T) and non-transmitted (NT) polygenic scores for educational attainment (EA) and ADHD on offspring’s academic achievement, ADHD symptoms at home and ADHD symptoms at school

|  | Model 1 |  |  | Model 2 |  |  | Model 3 |  |  |
| --- | --- | --- | --- | --- | --- | --- | --- | --- | --- |
|  | Beta | R<sup>2</sup> (%) | p | Beta | R<sup>2</sup> (%) | p | Beta | R<sup>2</sup> (%) | p |
| **Academic achievement (N = 1,120)** |  |  |  |  |  |  |  |  |  |
| EA<sub>T</sub> | .238 (.18;.30) | 5.7 | 7x10<sup>-15</sup> |  |  |  | .233 (.17;.30) | 5.4 | 2x10<sup>-13</sup> |
| EA<sub>NT</sub> | .034 (-.03;.10) | 0.1 | .284 |  |  |  | .033 (-.03;.10) | 0.1 | .316 |
| ADHD<sub>T</sub> |  |  |  | -.077 (-.14;-.01) | 0.6 | .022 | -.021 (-.09;.04) | 0.0 | .524 |
| ADHD<sub>NT</sub> |  |  |  | -.017 (-.08;.05) | 0.0 | .610 | -.006 (-.07;.06) | 0.0 | .857 |
| **ADHD symptoms at home (N = 2,518)** |  |  |  |  |  |  |  |  |  |
| EA<sub>T</sub> | -.125 (-.17;-.08) | 1.6 | 4x10<sup>-8</sup> |  |  |  | -.088 (-.14;-.04) | 0.8 | 2x10<sup>-5</sup> |
| EA<sub>NT</sub> | -.010 (-.05;.03) | 0.0 | .669 |  |  |  | -.017 (-.06;.03) | 0.0 | .488 |
| ADHD<sub>T</sub> |  |  |  | .165 (.12;.21) | 2.7 | 1x10<sup>-13</sup> | .141 (.10;.19) | 2.0 | 1x10<sup>-9</sup> |
| ADHD<sub>NT</sub> |  |  |  | -.030 (-.08;.02) | 0.1 | .190 | -.036 (-.08;.01) | 0.1 | .127 |
| **ADHD symptoms at school (N = 1,969)** |  |  |  |  |  |  |  |  |  |
| EA<sub>T</sub> | -.131 (-.18;-.08) | 1.7 | 1x10<sup>-7</sup> |  |  |  | -.104 (-.15;-.05) | 1.1 | 8x10<sup>-5</sup> |
| EA<sub>NT</sub> | -.012 (-.06;.04) | 0.0 | .637 |  |  |  | -.005 (-.06;.05) | 0.0 | .848 |
| ADHD<sub>T</sub> |  |  |  | .126 (.08;.18) | 1.6 | 3x10<sup>-7</sup> | .097 (.05;.15) | 0.9 | 2x10<sup>-4</sup> |
| ADHD<sub>NT</sub> |  |  |  | .029 (-.02;.08) | 0.1 | .243 | .024 (-.03;.07) | 0.1 | .353 |

**Figure 1.**
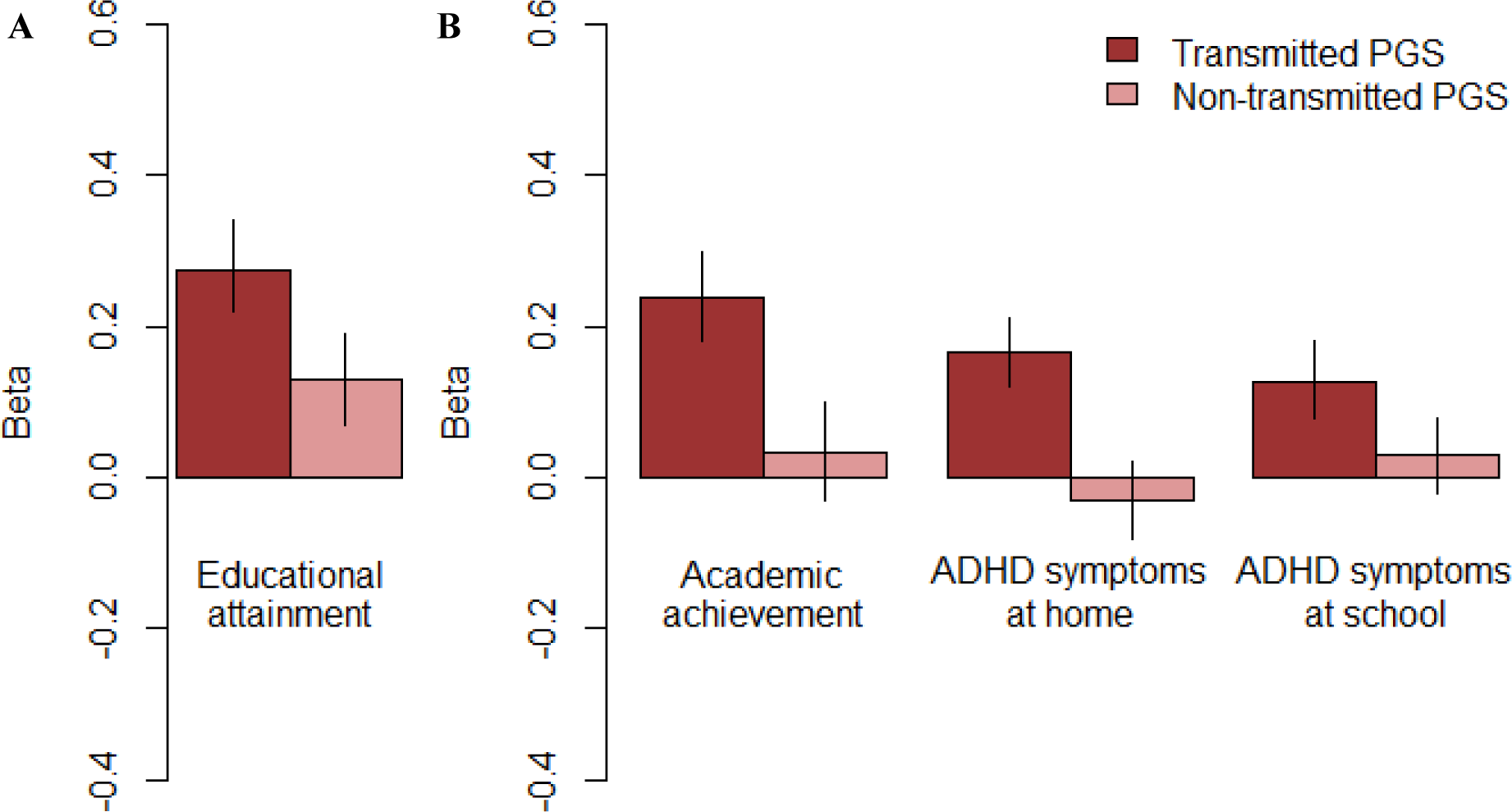
Effects (with 95% CI) of (A) the EA PGSs on educational attainment in adults and of (B) the EA PGSs on academic achievement in children and of the ADHD PGSs on childhood ADHD symptoms at home and at school.

**Figure 2.**
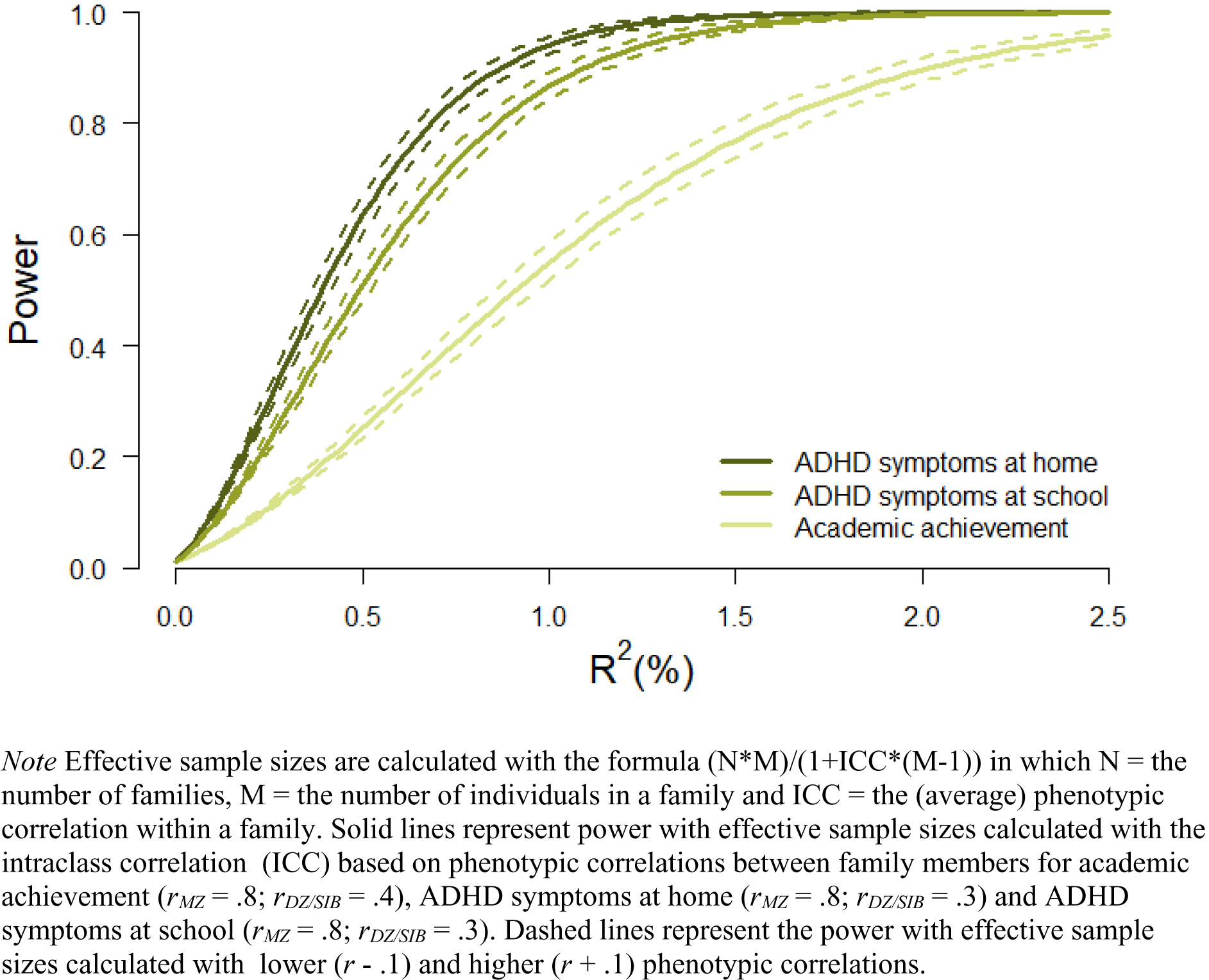
Power to detect the fixed effect of the non-transmitted PGS (expressed in R^2^) based on the calculated effective sample sizes for academic achievement (N_effective_ = 727), ADHD symptoms at home (N_effective_ = 1,702) and ADHD symptoms at school (N_effective_ = 1,352)

**Figure 3.**
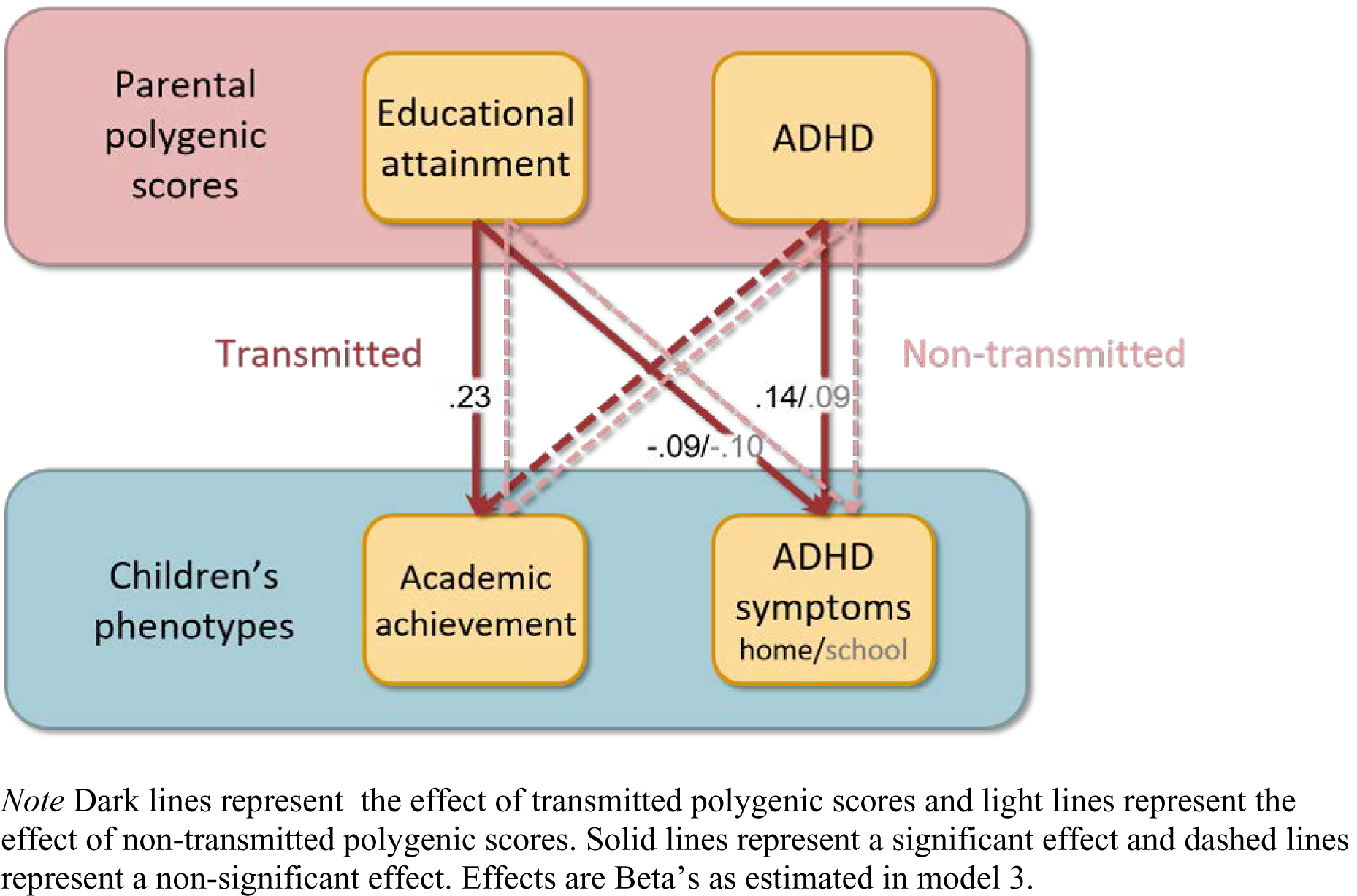
Schematic overview of the results

To determine if we had enough power to detect an effect of genetic nurturing on our childhood outcomes, we carried out a series of power analyses. Figure 2 displays the power, based on the effective sample sizes (i.e. the number of independent cases corresponding to the number of clustered cases), to detect a fixed effect of the non-transmitted PGS on each of the outcomes. The analyses indicated that there was sufficient power (.80) to detect an R^2^ explained by the non-transmitted PGS of 1.6%, 0.7% and 0.9% for, respectively, academic achievement, ADHD symptoms at home and ADHD symptoms at school.

## Discussion

In the current study we showed that both transmitted and non-transmitted alleles related to EA impact adult’s lifetime EA. Thus we replicate findings of other research groups (Bates et al. 2018; Kong et al. 2018) in an independent sample in a different cultural setting. This affirms that genetic nurturing, the effect of parental alleles over above the effects of transmitted alleles, influences adult EA, indicating that parents contribute to the offspring environment relevant to EA.

In childhood, in contrast, we found only effects of transmitted alleles, despite having sufficient power to pick up small effects of non-transmitted alleles. Given the disparity between the twin literature on academic achievement and ADHD, this is in line with our hypothesis for ADHD, but not for academic achievement. The effect of genetic nurturing in the current study was non-significant, and close to zero (R^2^ ∼ 0.1%). Regarding academic achievement, the twin literature shows a small but significant effect of common environment in the Netherlands (de Zeeuw et al. 2016), so we had expected to find a small but significant effect of genetic nurturing. However, environmental transmission was not found for the resemblance in reading achievement between parents and their children (Swagerman et al. 2017). It could be that the estimated common environment effect includes environmental influences shared between siblings that are independent of parent genes, indicating a vertical, not horizontal, environmental transmission effect.

How can we reconcile the effect of genetic nurturing on education being absent in childhood (R^2^ of 0.1%) but present in adulthood (R^2^ of 1.7%)? First, academic achievement scores at age 12 and the highest obtained degree in adulthood are different traits. We used PGSs for EA, so the trait in adulthood. Hence, the effect of the transmitted alleles is expected to be somewhat higher for education in adulthood (7.6%) than childhood (5.7%). Nevertheless, the effects differ much more for the non-transmitted alleles, so this does not seem to explain the full story. An increasing genetic nurturing effect is consistent with a slightly increasing common environment during secondary school (Rimfeld et al. 2018) and a substantial common environment in adults’ EA (Branigan et al. 2017).

We speculate that the common environmental effect (and genetic nurturing effect) increases in secondary school due to educational tracking. Tracking in the Netherlands takes place from age 12 onwards, partly based on the academic achievement test that we analysed here. Parents’ EA still predicts test scores after accounting for genetic influences (de Zeeuw et al. 2019). Moreover, children from higher-educated parents are more likely to enrol in and complete a higher track than expected based on the test result (van Spijker et al. 2017). Together, parents’ EA (which may induce genetic nurturing) seems to play a role in (Dutch) secondary school beyond children’s inherited abilities. Next, due to tracking, the school environment that children experience in secondary school varies much more than in primary school. More environmental variance would increase the contribution of the common environment in explaining achievement differences. In addition, due to tracking, children would experience a school level that broadly aligns with their genetic potential, thereby inducing a gene-environment correlation. In the classical twin design with uncorrelated genetic and common environmental factors, the correlation between these factors will show up as common environment variance.

No genetic nurturing in childhood was expected for ADHD symptoms. It was expected based on the lack of common environment in the ADHD literature (Faraone and Larsson 2019) and the large contribution of dominant genetic effects (Rietveld et al. 2003; Nikolas and Burt 2010). Our confidence in the null-result for AP is strengthened by the fact we had sufficient power (.80) to detect relatively small effects (at home: 0.7% and at school 0.9%) and by fact that we observe it for both children’s home and school situation. Apparently, the aspects of the home environment that are associated with parents genetic liability for ADHD do not impact children’s cognitive and behavioural development.

The absence or presence of the effect of non-transmitted alleles is interesting, but so is the ratio between non-transmitted and transmitted. As all parental alleles induce genetic nurturing, those transmitted capture both genetic nurturing and a direct genetic effect (Kong et al. 2018). The direct genetic effect is thus the difference between effects of transmitted and non-transmitted PGSs. Kong et al. (2018) found for EA that the effect of genetic nature was almost half the size of the direct genetic effect. Here we found for EA that genetic nurturing was almost as large as the direct genetic effect. This ratio falls in the range of ratios for other traits in adulthood that Kong et al. (2018) report. In childhood, however, we found that the ratio between genetic nurturing and direct genetic was only a sixth for achievement and a tenth for ADHD symptoms. Thus, the effect of genetic nurturing that we found in childhood was not only tiny in itself, but also minor relative to the PGS’s estimated direct genetic effect.

We set out to test investigate transmission both within and between domains. Only effects of transmitted PGSs were significant, so all associations were attributable to genetic transmission, and not genetic nurturing. Children’s ADHD symptoms were predicted by both PGSs based on transmitted alleles associated with ADHD and EA. This is consistent with the genetic correlation between academic achievement and ADHD (e.g. Liu et al. 2019). Yet, academic achievement was only influenced by parents’ EA associated alleles. In light of the current results, earlier reported associations between household chaos and academic achievement (Johnson et al. 2009; Hanscombe et al. 2011) seem not environmentally, but genetically transmitted via education-related alleles. However, this is not in line with a British twin study in which it was concluded that chaotic homes predict poor academic achievement due to a combination of shared genetic and environmental effects (Hanscombe et al. 2011). These seemingly conflicting conclusions may be due to population differences: the Dutch educational system is egalitarian, resulting in reduced effects of the common environment (Kovas et al. 2013; de Zeeuw et al. 2016).

A significant effect of the non-transmitted PGS evidences that parenting relating to EA matters. However, the absence of an effect of the non-transmitted PGS does not necessarily mean that these associated parenting behaviours do not matter. It is possible that the link between the education related parenting behaviours and childhood academic achievement, might go directly through transmission of genetic effects, but that the effects of parenting behaviours not associated with EA go through the environment, or in other words, genetic nurturing. These parental behaviours could be shaped by other parental traits, more specifically, parental traits with a low genetic correlation with EA, for example personality, well-being, and health (Bulik-Sullivan et al. 2015). Adding non-transmitted PGSs for a large range of different parental characteristics may help characterize a child’s environment and to pinpoint the parental behaviours that do have an impact on offspring development.

It was recently demonstrated that passive gene-environment correlations contribute to the predictive power of PGSs for cognitive traits but not for ADHD. Selzam et al. (2019) leveraged the fact that within-family analyses account for passive gene-environment correlations. In contrasting within- and between-family analyses, it was shown that PGS prediction estimates for cognitive traits were greater between families than within families. In contrast, within- and between-family estimates were similar for the prediction of ADHD. Passive gene-environment correlations are picked up in our design by genetic nurturing. Therefore, their conclusions are compatible with our findings of genetic nurturing for EA but not ADHD.

The use of PGSs in the genetic nurturing design means that all GWAS limitations carry over to the current study, including that the summary statistics can be affected by remaining population stratification, the use of alternative phenotype definitions in discovery GWAS samples, and a different population background as the Dutch population used here (De La Vega and Bustamante 2018). Nonetheless, the genetic nurturing design comes with unique strengths. It can demonstrate influences of the family environment without relying on self-reports. Studies on, for example, the impact of growing up in a chaotic household typically rely on parents’ own judgement of their home situation or of their own ADHD symptoms. Here we used unbiased ADHD and EA related genetic variation as a tool to demonstrate causal environmental effects un-confounded by genetic effects. This tool requires for the child generation genotypes and phenotypes, but for the parent generation only genotypes. Nevertheless, adding parental phenotypes opens possibilities to further disentangle effects of direct genetic liabilities, causal environments, and forms of gene-environment correlation. Establishing genetic nurturing can then be followed-up by pinpointing its mechanism, for example, whether it affects children via cognitive stimulation or household chaos (Wertz 2019).

In the current study, we found a genetic effect mediated by the environment (‘genetic nurturing’) on EA, in addition to the direct genetic effect. The genetic nurture method tests influences of parenting behaviour beyond gene-environment correlation. Not all direct genetic effects are captured and therefore not all gene-environment correlations are captured. But regardless of the proportion of the total genetic effect captured, we should bear in mind that the child does not have non-transmitted alleles, so effects of non-transmitted alleles always evidence causal effects via the environment. For academic achievement and behaviour in childhood, we only found evidence for direct genetic effects of EA and ADHD. This suggests that there is no environmental transmission of parental genetic effects and they are solely transmitted via genetic transmission. As this method is new to the field and still in development, replication of our findings is warranted, preferably with larger samples and more powerful PGSs. Nevertheless, from the current findings we conclude that reported associations between home characteristics related to parental EA and ADHD and child outcomes seem to be mainly a marker of genetic effects shared by parents and children.

## Acknowledgements

We are grateful to the twin families and the teachers for their participation. We gratefully acknowledge research program ‘Consortium on Individual Development (CID)’ which is funded through the Gravitation program of the Dutch Ministry of Education, Culture and Science and the Netherlands Organization for Scientific Research (NWO: 0240-001-003); ‘Decoding the gene-environment interplay of reading ability’ (NWO: 451-15-017); ‘Netherlands Twin Registry Repository: researching the interplay between genome and environment’ (NWO: 480-15-001/674); ‘Twin-family study of individual differences in school achievement’ (NWO: 056-32-010); ‘Longitudinal data collection from teachers of Dutch twins and their siblings’ (NWO: 481-08-011); ‘KNAW Academy Professor Award (PAH/6635)’.

